# Carbonized Tetracycline: a new class of nanomaterial with tuneable antioxidant, reduced cytotoxicity, immunomodulatory, and osteogenic properties

**DOI:** 10.1101/2023.10.30.564661

**Authors:** Saptarshi Mandal, Ramkumar T Annamalai

**Affiliations:** Department of Biomedical Engineering, University of Kentucky, Lexington, KY 40506; Department of Biomedical Engineering, University at Buffalo, Buffalo, NY 14260

**Keywords:** Carbon dot, Graphene quantum dot, Macrophage polarization, Osteogenesis, Metalloproteinase

## Abstract

Tetracycline (TET), a broad-spectrum antibiotic, also possesses different non-antibiotic activities such as inhibition of metalloproteinase (MMP), anti-inflammatory, antioxidant, high bone affinity, etc. However, the comparatively low efficacy of these non-antibiotic properties along with adverse effects such as hyperpigmentation, phototoxicity, long-term skeletal retention, etc. have not helped their broad utilization similar to their use as an antibiotic. In a unique attempt to improve the non-antibiotic properties while reducing the adverse effects, we converted the TET to nano-carbons through partial carbonization. After sorting out two water-dispersible C-TETs (C-TET_HT_ - hydrothermal and C-TET_HP_ - hot plate) based on their improved antioxidant activity, they have been characterized through a host of analytical techniques that showed distinct differences in morphology, size, shape, and surface functionality. Excitingly, the C-TET_HT_ and C-TET_HP_ have shown differential biological activity in a dosage and time-dependent manner in terms of cytotoxicity, immunomodulation, and osteogenic activity that was found to be associated with their carbonized parameters. Overall, the carbonized nano-drugs, C-TET_HT_ and C-TET_HP_ have presented substantial early promises on their non-antibiotic properties that could be further explored to develop into some effective therapeutics.

## 1. Introduction

Carbonization is the process of converting organic matter through heating. In the last few decades, carbonized nanomaterials have gained substantial attention in several areas of biomedical engineering, especially nanotherapeutics, due to large surface-to-volume ratio, facile functionalization, tuneable optical properties, etc.^1, 2^ Among these carbon-based nanomaterials, the tiny carbon dots have possessed additional advantages in terms of easy and inexpensive synthesis, decent water dispersibility, good biocompatibility, etc.^3, 4^ These tiny nano-carbons could be synthesized by partially carbonizing any organic matters through hydrothermal or solvothermal pyrolysis, hot plate or microwave heating, etc.^3^ Notably, it was reported that a few properties of the starting raw materials could be retained after carbonization also.^3, 5^ For example, the carbonized form of a few small drug molecules, such as aspirin^6^ levofloxacin,^7^ 2-methoxy-1, 4-naphthoquinone,^8^ and ciprofloxacin,^9^ are shown to have better therapeutic properties. Although the mechanisms for the enhanced therapeutic properties are not fully clear, the change in water dispersibility, systemic circulation half-life, cellular uptake, and retention of the carbonized materials are thought to play a crucial role. The carbonized form of a few small drug molecules has also shown efficacy in diseases other than the traditional use of that drug. For example, metformin, a first-choice drug for type 2 diabetes, has shown good results in its carbonized form in periodontitis and colorectal cancer model.^10, 11^ Inspired by these revelations, herein, we have made an effort to partially carbonize a drug Tetracycline (TET) to improve its non-antibiotic properties.

Tetracyclines are a broad-spectrum class of antibiotics that have grabbed decent research attention for their various non-antibiotic properties such as inhibition of metalloproteinase (MMP) activity, anti-inflammatory activity, antioxidant effect, high affinity for bone, etc. making tetracyclines drugs with colossal therapeutic potential.^12–14^ They inhibit MMP activity by chelating metal ions required for the activity of MMP,^15^ show antioxidant effect by scavenging the reactive oxygen species (ROS) through its phenolic ring,^16^ and are attracted towards bone because of their affinity for calcium.^13^ Tetracyclines and osteogenesis is a half-a-century-old research topic that is yet to be clarified.^17^ They show anti-inflammatory effects by inhibiting the production of pro-inflammatory cytokines, leukocyte migration, and inducible nitric oxide synthase activity.^18^ Based on these properties Doxycycline, Minocycline, and TET have been investigated to treat various bone diseases such as osteoarthritis, osteopenia, etc.^13, 19^ Efforts were also made in developing chemically modified tetracyclines (CMTs) that have no antibiotic effect but have similar or enhanced non-antibiotic properties.^20, 21^ However, the use of tetracyclines has been hindered due to comparatively low efficacy, and various adverse effects like phototoxicity, skeletal retention, hyperpigmentation, etc.^13, 22^ Hence, a need for research focusing on reducing the toxicity of tetracyclines while enhancing the efficacy of non-antibiotic properties is substantial.

Carbon nanomaterials have shown good initial promises in therapies critical to bone defects and related diseases, based on their good mechanical properties, large surface area, tuneable surface characteristics, efficient osteogenic differentiation, etc.^23^ They have shown immunomodulatory properties through changing cytokine production, altering immune cell maturation, etc.^24^ Nano-carbons such as carbon dots have been reported to act as an antioxidant, while they can act as an oxidant in the presence of light.^25^ A dexamethasone-derived carbon dot was also reported for its osteogenic and immunomodulatory activity,^26^ while a folic acid-derived CD protected the chondrocytes.^27^ However, systemic investigation on how various parameters of nanocarbons such as surface morphology, surface charge, size, shape, etc, could affect the cell fates is lacking specifically.^23^ Herein, we endeavored to synthesize the carbonized nanomaterials with different surface morphology, size, and shape so that we could shed light on their intercorrelations with the cell fates.

Among the tetracyclines class of antibiotics, we have chosen the TET, the first generation tetracyclines, and a natural compound to partially pyrolyze through hydrothermal and hot plate heating techniques. The synthesis of the carbonized TET (C-TET) was optimized by varying different pyrolytic conditions such as temperature, pH, pyrolysis time, and method. of pyrolysis. Among the five synthesized C-TETs, we have picked out two C-TETs based on their antioxidant activity. Interestingly, these two C-TETs appeared as hydrothermally pyrolyzed and a hot plate heated C-TETs and named C-TET_HT_ and C-TET_HP_. The two C-TETs were then extensively characterized through a host of standard analytical techniques. The cytotoxicity of the C-TETs was assessed through cell metabolism and proliferation assay and compared with TET. How the C-TET_HT_ and C-TET_HP_ could polarize the macrophage phenotypes was evaluated. Whether C-TET_HT_ and C-TET_HP_ can promote chondrogenic differentiation in an inflammatory condition was also tested. The osteogenic potential of the C-TETs was then assessed. From the cytotoxicity to the macrophage polarization, chondroprotective, and osteogenesis, the C-TET_HT_ and C-TET_HP_ demonstrated differential results. Overall, our attempt to synthesize and evaluate TET-based carbonized nano-drugs (C-TETs) have shown substantial promise for further studies in musculoskeletal diseases with tuneable antioxidant, reduced cytotoxicity, immunomodulatory, and osteogenic properties.

## 2. Materials and Methods

### 2.1. Synthesis of C-TET

C-TETs were synthesized either by open pyrolysis on a hot plate or through hydrothermal treatment. Briefly, 0.5 mmol of TET (Millipore Sigma) was dispersed or dissolved in 5 mL of water, 1.5% HCl solution, 5% H_3_PO_4_ solution, or 0.2M NaOH solution. Then the dispersion or solution of TET was transferred to a 25 mL hydrothermal reaction vessel. The hydrothermal autoclave was heated inside a hot air oven for some specific time and temperature. After cooling down to room temperature the crude carbonized mixture dispersion was neutralized either with dilute HCl or dilute NaOH solution. Then it was centrifuged at rpm 4K for 10 min and the supernatant was collected. The supernatant was dialyzed with a membrane of MWCO 2KDa (Millipore Sigma) for 16 hours with occasional changes of water. After that, the retentate was passed through a 0.22 µm nylon syringe filter (VWR). Finally, the purified C-TETs were collected and stored for further use. For the hot plate pyrolysis method, the TET dispersion or solution was taken in a 125 mL Erlenmeyer flask and then heated on a hot plate under a fume hood for a specific time and temperature. Water was added occasionally to prevent complete evaporation of the solvent (water). The rest of the synthesis and purification procedures are similar to that of the hydrothermally prepared C-TETs.

### 2.2. Antioxidant activity (DPPH assay)

The antioxidant activity of the samples was determined by 2,2-Diphenyl-1-picrylhydrazyl (DPPH) radical (Millipore Sigma) scavenging activity of the samples.^28,29^ Briefly, the C-TETs dispersion or TET solution of different concentrations was prepared in 100 mM Tris.HCl buffer, pH 7.4. DPPH solution (100 µg/mL) in ethanol was freshly prepared. In a 96-well plate, 100 µL DPPH solution was added to the 100 µL of different concentrations of samples. 100 µL DPPH solution in 100 µL Tris.HCl buffer was used as blank, whereas 100 µL ethanol in 100 µL of each of the samples was considered as control. The well plate was then incubated in the dark for 30 minutes, and the absorbance was measured at 517 nm using a Biotek PowerWave HT microplate reader. The radical scavenging activity (RSA) of the samples was calculated as: 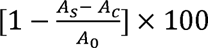. A_s_ is the absorbance of the sample, A_c_ is the absorbance of the control and A_0_ is the absorbance of the blank. The IC_50_ (sample concentration for 50% inhibition of the DPPH radical) was calculated from the calibration curve in the linear range by plotting the sample concentration and corresponding RSA value.

### 2.3. Characterization of C-TET

An FEI Talos F200X (Thermo Scientific) transmission electron microscope (TEM) was used for imaging, selected area electron diffraction (SAED), and energy dispersive X-ray spectroscopy (EDS) studies. Diluted samples at a concentration of 100 µg/mL (water as diluent) were drop cast on formvar film gold 100 square mesh TEM grids and air dried. Dynamic light scattering (DLS) and zeta potential measurements of diluted samples in 1x PBS at pH 7.4 were performed on a Zetasizer Nano ZS (Malvern Panalytical). The powder X-ray diffraction (pXRD) study was performed at room temperature on a Bruker-AXS D8 DISCOVER diffractometer using Cu-Kα as the X-ray source (λ = 1.54Å). The dried samples were put on the sample holder and scanned at 2θ angle from 5° to 60° with a scan step of 0.02°. Raman spectra of the samples were obtained using a DXR Raman Microscope (Thermo Scientific) at an excitation wavelength of 532 nm. The concentrated samples from the stock were drop-cast on a glass slide and air-dried completely before collecting the Raman spectra. X-ray photoelectron spectroscopy (XPS) was performed on a Thermo Scientific K-Alpha XPS system where the dried samples were directly mounted on the sample holder. ^1^H NMR spectra were obtained on a Varian 500 MHz NMR instrument (VNMRS500) using D_2_O as the solvent. FTIR…. Dried samples were pelletized with KBr (Thermo Scientific Chemicals) and scanned over a wavenumber range of 4000 cm^-1^ to 500 cm^-1^. Diluted samples in 1x PBS pH 7.4 at a concentration of 10 µg/mL were taken in a quartz cuvette of pathlength 1 cm and scanned over a wavelength range of 800 nm to 200 nm on an Evolution^TM^ 201 (Thermo Scientific) UV-Vis spectrophotometer. Then the same samples were scanned on a Fluoromax Plus-C (Horiba Scientific) steady-state spectrofluorometer at different excitation wavelengths keeping the other parameters the same.

### 2.4. Photoinduced ROS generation (DHR 123 assay)

10 µL of 10 µM dihydrorhodamine (DHR) 123 (Millipore Sigma) solution was added to 190 µL of C-TET dispersion in 1x phosphate-buffered saline (PBS), pH 7.4 of C-TETs concentration 10 µg/mL and 100 µg/mL. DHR 123 solution (10 µL, 10 µM) in 1x PBS buffer (190 µL, pH 7.4) without any C-TET was used as the control. All the samples were irradiated with UV light (wavelength 365 nm) for 30 minutes. 100 µL of each sample was then transferred to a black 96 well plate and the fluorescence was measured at an excitation/emission wavelength of 488/530 nm using a SpectraMax Gemini XS microplate spectrofluorometer (Molecular Devices, CA).^30^

### 2.5. Cell culture

Normal human lung fibroblast (NHLF) cell line (Lonza, CC-2512) was used to study the cytotoxicity of the C-TET_HT_ and C-TET_HP_. NHLF was grown in Dulbecco’s modified eagle medium (DMEM, high glucose, GlutaMAX, Gibco) supplemented with 10% fetal bovine serum albumin (FBS, ThermoFisher), 1% Antibioic-Antimycotic (AA, Gibco) and 0.2% Normocin. The cells were seeded in 24-well plates at 25,000 cells/well. After 24 hr of incubation, the cells were treated with 10 µg/mL and 100 µg/mL of TET solution and C-TET_HT_ and C-TET_HP_ dispersions in media. Cells without TET or C-TETs treatment were used as the control.

IC-21 macrophage cell line (ATCC, TBI-186) developed from C57BL/6 murine background was used for macrophage polarization studies. The cells were cultured in Roswell Park Memorial Institute (RPMI) 1640 medium (ATCC modification, Gibco) supplemented with 10% FBS, and 1% AA. The cells were seeded in 12-well plates at 80,000 cells/well and incubated for 24 hr. To polarize the macrophages from M0 to M1, the 24 hr incubated cells were treated with 25 ng/mL interferon-gamma (IFN-γ) and 50 ng/mL lipopolysaccharide (LPS) and incubated for another 24 hr. The M0 and M1 macrophages were then treated with 10 µg/mL and 100 µg/mL of the C-TET_HT_ and C-TET_HP_ dispersions in media. Cells without C-TETs treatment were used as the control.

The chondrogenic cell line ATDC5 (Millipore Sigma) was cultured in DMEM/Ham′s F12 (1:1) media supplemented with 2mM Glutamine, 5% FBS, 1% AA, and 0.2% Normocin. The cells were seeded in 12-well plates at 80,000 cells/well and incubated for 48 hr. Then the cells were treated with 10 ng/mL IL-1b. After 24 hr of incubation, cells were treated with 10 ng/mL and 100 µg/mL of the C-TET_HT_ and C-TET_HP_ dispersions in media containing 10 ng/mL IL-1b. Cells without any C-TETs treatment and cells without IL-1b treatment were also cultured for control.

The osteoblast cell line MC3T3-E1 (RCB1126, RIKEN) was used to study the osteogenic potential of the C-TET_HT_ and C-TET_HP_. The cells were grown in Minimum Essential Medium α (MEM α, nucleosides, no ascorbic acid, Gibco) media supplemented with 10% FBS, 1% AA, and 0.2% Normocin. The cells were seeded both in normal media and osteogenic media in 24-well plates at 40,000 cells/well. The osteogenic media contains 10 mM β-glycerolphosphate (Millipore Sigma) and 0.2 mM L-ascorbic acid-2-phosphate (Millipore Sigma) along with all the contains of normal media. After 48 hr of incubation, the cells were treated with 5 µg/mL and 25 µg/mL of the C-TET_HT_ and C-TET_HP_ dispersions in media. Cells without any C-TETs treatment were considered as the control.

All the cells were grown at 37 °C temperature and maintained at 5% CO_2_ inside an incubator. The media was changed every 2 days.

### 2.6. PrestoBlue assay

The metabolic activity of the cells was quantified through PrestoBlue (Invitrogen) assay at specified time points.^31^ Cells were incubated at 37 ℃ for 1 hr with 1x PrestoBlue solution containing the fresh media. 200 µL from each well was transferred to a black 96-well plate and the fluorescence was measured at an excitation/emission wavelength of 560/590 nm on SpectraMax Gemini XS microplate spectrofluorometer (Molecular Devices, CA).

### 2.7. Cell viability

The cell viability was determined by LIVE/DEAD™ Viability/Cytotoxicity Kit (Invitrogen) at specified time points.^32^ Briefly, cells were washed with 1x PBS buffer. To each well 200 µL of the assay solution containing 2 µM of calcein AM and 4 µM of ethidium homodimer-1 in 1x PBS was added and incubated at room temperature for 30 min. The cells were then imaged on a fluorescence microscope. The fluorescence was then quantified using ImageJ software and plotted accordingly.

### 2.8. Quantitative gene expression

Macrophage polarization, chondrogenesis in an inflammatory condition, and osteogenic potential of the C-TET_HT_ and C-TET_HP_ were studied with quantitative gene expression analysis. The samples were collected in TRIzol reagent (Invitrogen) and the RNA was isolated according to the manufacturer’s protocol. The RNA was quantified using a NanoDrop One spectrophotometer (Thermo Scientific). SuperScript III Platinum One-Step qRT-PCR Kit (ThermoFisher) and TaqMan probes (ThermoFisher) were used for the gene expression study using a QuantStudio 3 real-time PCR system (ThermoFisher). The following TaqMan probes were used for the studies: IL6 (Mm00446190_m1), NOS2 (Mm00440502_m1), TNF-α (Mm00443260_g1), IL-1b (Mm00434228_m1), IL10 (Mm00439616_m1), ARG1 (Mm00475988_m1), TGF-β (Mm01178820_m1), SOX9 (Mm00448840_m1), ACAN (Mm00545794_m1), COL2A1 (Mm01309565_m1), COX2 (Mm03294838_g1), ADAMTS5 (Mm00478620_m1), ADAMTS4 (Mm00556068_m1), MMP3 (Mm00440295_m1), MMP1a (Mm00473485_m1), MMP13 (Mm00439491_m1), MMP9 (Mm00442991_m1), RUNX2 (Mm00501584_m1), COL1A1 (Mm00801666_g1), ALP1 (Mm00475834_m1), BGLAP (Mm03413826_mH), SPP1 (Mm00436767_m1), IBSP (Mm00492555_m1), SP7 (Mm04209856_m1). GAPDH (Mm99999915_g1) was used as housekeeping control. The gene expression data were presented as heatmaps.

### 2.9. ALP activity assay

Alkaline phosphatase (ALP) activity was measured fluorometrically using a commercially available kit (BioVision) according to the manufacturer’s protocol.^32^ Briefly, the media was removed from the cell wells and washed with 1x PBS. 100 µL of the assay buffer was added to each well. the cells were homogenized through sonication and then centrifuged at 13,000g for 3 min. In a black 96-well plate 100 µL of the assay buffer was added, followed by 10 µL of the supernatant, and then mixed with 20 µL of the 0.5 mM 4-Methylumbelliferyl phosphate disodium salt (MUP) substrate solution. After incubating the plate for 30 min in the dark 20 µL stop solution was added to the mixed solution. The fluorescence was measured at an excitation/emission wavelength of 360/440 nm on a SpectraMax Gemini XS microplate spectrofluorometer. The ALP activity was measured from the standard curve. The standard curve was generated by using different amounts of MUP substrate (0 – 0.5 nmol/well) with the supplied ALP enzyme solution.

### 2.10. Alizarin red staining

1% (w/v) alizarin red solution of pH 4.2 was prepared by dissolving alizarin red powder (Millipore Sigma) in water and then adjusting the pH accordingly with dilute HCl (Acros Organics) solution. After aspirating the growth media, the cells were washed with 1x PBS and fixed in a 4% buffered formalin solution at room temperature for 60 min. Then the formalin solution was removed, and the cells were stained with 1mL of 1% alizarin red solution per well for 10 min. The alizarin red solution was then removed, the cells were washed thrice with distilled water, and imaged using bright-field microscopy. The cells were then destained with 0.3 mL of 5% sodium dodecyl sulfate (Millipore Sigma) solution in 0.5 N HCl for 15 min. The absorbance of the destained extracted solution was measured at wavelength 405 nm using a BioTek PowerWave HT microplate spectrophotometer.

### 2.11. Statistics

All measurements were performed with at least three replicates. Data are plotted as either means with error bars representing the standard deviation. One-way ANOVA was used to identify significance within the groups and Holm-Sidak pairwise comparison was followed to identify the significant groups. All statistics tests were done using SigmaPlot. Differences with a *P-*value < 0.05 were considered statistically significant.

## 3. Results and Discussion

### 3.1. Synthesis of C-TETs

C-TETs were synthesized following two well-established methods for the synthesis of carbon dots – hydrothermal and direct hot plate pyrolysis (**Table 1**, **Figure 1A**).^3^ Our initial efforts of synthesizing C-TET in neutral water and dilute acidic solution ended up with large black precipitates. C-TETs were then synthesized in 0.2M NaOH, and we varied the pyrolysis time and temperature to optimize the yield. Hydrothermal pyrolysis at 150℃ (yield ∼ 82%) was found better in terms of yield of C-TETs than pyrolyzing at 180℃ (yield ∼52%), whereas pyrolyzing for a prolonged time (15 hours as compared to 6 hours) at 150℃ further increased the yield (∼88%). Quite notably, the hot plate method of pyrolysis provides a yield of ∼62%. Purification of the C-TETs was performed in three steps, centrifugation, dialysis, and then followed by syringe filtration. Centrifugation was performed to remove the large-carbonized materials from the crude pyrolyzed mixture. The dialysis was done to eliminate any molecular byproducts and unreacted raw materials (if any were present). Syringe filtration of the retentate further eliminates aggregates providing well-dispersed C-TETs for further studies. Overall, the synthesis yields of C-TETs demonstrate the superiority of the base-catalyzed pyrolytic treatment over the acid-catalyzed or neutral pyrolytic treatments. On another hand, whereas TET has poor water solubility, the successfully synthesized C-TETs possess good water dispersibility.

**Figure 1.**
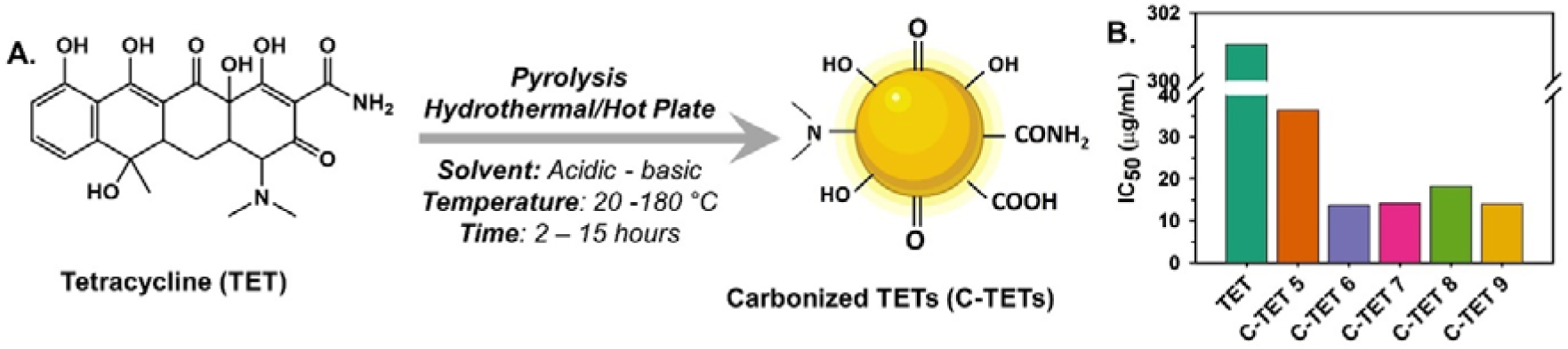
**A.** Synthesis parameters for C-TETs. **B.** Antioxidant activity (IC_50_) of TET and C-TETs showed a great increase in radical scavenging activity after partial carbonization.

**Table 1.** Various conditions for the synthesis of C-TETs.

| C-TETs | Solvent, mL | Temp., °C | Time, h | Method | Outcome |
| --- | --- | --- | --- | --- | --- |
| C-TET 1 | Water, 5 mL | 180 | 6 | Hydrothermal | Large black precipitate inside the reaction vessel |
| C-TET 2 | Water, 5 mL | 150 | 6 | Hot plate | Black precipitate, poor water dispersibility |
| C-TET 3 | 1.5% HCl solution, 5 mL | 150 | 6 | Hydrothermal | Large black precipitate inside the reaction vessel |
| C-TET 4 | 5% H<sub>3</sub>PO<sub>4</sub> solution, 5 mL | 150 | 4 | Hydrothermal | Large black precipitate inside the reaction vessel |
| C-TET 5 | 5% H<sub>3</sub>PO<sub>4</sub> solution, 5 mL | 150 | 6 | Hot plate | Yield ~62% |
| C-TET 6 | 0.2 M NaOH solution, 5 mL | 150 | 6 | Hydrothermal | Yield ~82% |
| C-TET 7 | 0.2 M NaOH solution, 5 mL | 180 | 6 | Hydrothermal | Yield ~52% |
| C-TET 8 | 0.2 M NaOH solution, 5 mL | 150 | 15 | Hydrothermal | Yield ~88% |
| C-TET 9 | 0.2 M NaOH solution, 5 mL | 150 | 6 | Hot plate | Yield ~61% |

### 3.2. Antioxidant activity of the C-TETs

After synthesizing several C-TETs, we had to select one or two C-TETs for further evaluation of their biological potential related to the non-antibiotic properties of TET. Among various non-antibiotic properties of TET, its radical scavenging activity or antioxidant property is relatively inexpensive and less time-consuming to evaluate. Herein, we assessed the antioxidant potential of all the C-TETs with synthesis yield >50% through DPPH assay and expressed as IC_50_ value, i.e., the amount of the sample required to scavenge 50% of the DPPH radical used in the assay (Figure 1B, Figure S1, Table S1). Remarkably, the antioxidant capacity of all the C-TETs was found 10-20 times better than that of TET. On the other hand, C-TETs synthesized at basic pH (NaOH) appeared 2-2.5 times better than the C-TET synthesized at acidic pH (H_3_PO_4_). Overall, C-TET 6, a C-TET synthesized by hydrothermal method, possesses the lowest IC_50_ value (13.81) followed by C-TET 9, a C-TET prepared by hot plate method (14.08). Hydrothermal treatment at a higher temperature or for a longer duration results in increased IC_50_ value (14.22 for C-TET 7 and 18.40 for C-TET 8, respectively) or decreased antioxidant activity. Based on the %yield and antioxidant capacity of the C-TETs, we picked C-TET 6 and C-TET 9 for further studies and named C-TET_HT_ and C-TET_HP_, respectively for easy understanding and comparison. As the main difference between the C-TET_HT_ and C-TET_HP_ is the method of pyrolysis, it could be a good exercise to know how the carbonization method could impact the therapeutic potential of carbonized drugs.

### 3.3. Characterization of C-TETs

C-TET_HT_ and C-TET_HP_ were characterized with a host of analytical techniques. TEM image analysis of C-TET_HT_ (**Figure 2A**) showed quasi-spherical morphology with an average size of 70±26 nm (Calculated with ImageJ software), whereas the TEM image of C-TET_HP_ (**Figure 2B**) showed no definable structure. Interestingly, in high-resolution TEM (HR-TEM) study, both the C-TET_HT_ and C-TET_HP_ revealed lattice fringes embedded in an amorphous carbon matrix (**Figure 2C-D, Figure S2**) and the corresponding fast Fourier transform (FFT) signifies amorphous nature with partial crystallinity of the C-TET_HT_ and the C-TET_HP_ (**Figure S2**).^33, 34^ pXRD showed weak and broad diffraction peak centered at ∼17° indicating partial graphitization mostly of disordered carbon with layer spacing ∼5.15 Å for both the C-TETs (**Figure 2E**).^33^ Notably the peak intensity for the C-TET_HP_ was quite low that suggests either poor graphitization or presence of tiny (<10 nm) carbon fragments.^34^ Raman spectroscopy was performed to understand the graphitization of the C-TETs (**Figure 2F**). Interestingly, any significant difference in the intensity ratio of D and G-band (I_D_/I_G_) for the C-TET_HT_ and C-TET_HP_ was not observed. Rather the second-order higher frequency bands 2D and D+G were found more intense and defined better for C-TET_HT_ than that for C-TET_HP_, indicating a more ordered and larger graphitic framework of C-TET_HT_ that correlates well with HR-TEM observations.^35^ ^33, 34^ pXRD showed weak and broad diffraction peak centered at ∼17° indicating partial graphitization mostly of disordered carbon with layer spacing ∼5.15 Å for both the C-TETs (**Figure 2E**).^33^ Notably the peak intensity for the C-TET_HP_ was quite low that suggests either poor graphitization or presence of tiny (<10 nm) carbon fragments.^34^ Raman spectroscopy was performed to understand the graphitization of the C-TETs (**Figure 2F**). Interestingly, any significant difference in the intensity ratio of D and G-band (I_D_/I_G_) for the C-TET_HT_ and C-TET_HP_ was not observed. Rather the second-order higher frequency bands 2D and D+G were found more intense and defined better for C-TET_HT_ than that for C-TET_HP_, indicating a more ordered and larger graphitic framework of C-TET_HT_ that correlates well with HR-TEM observations.^35^

**Figure 2.**
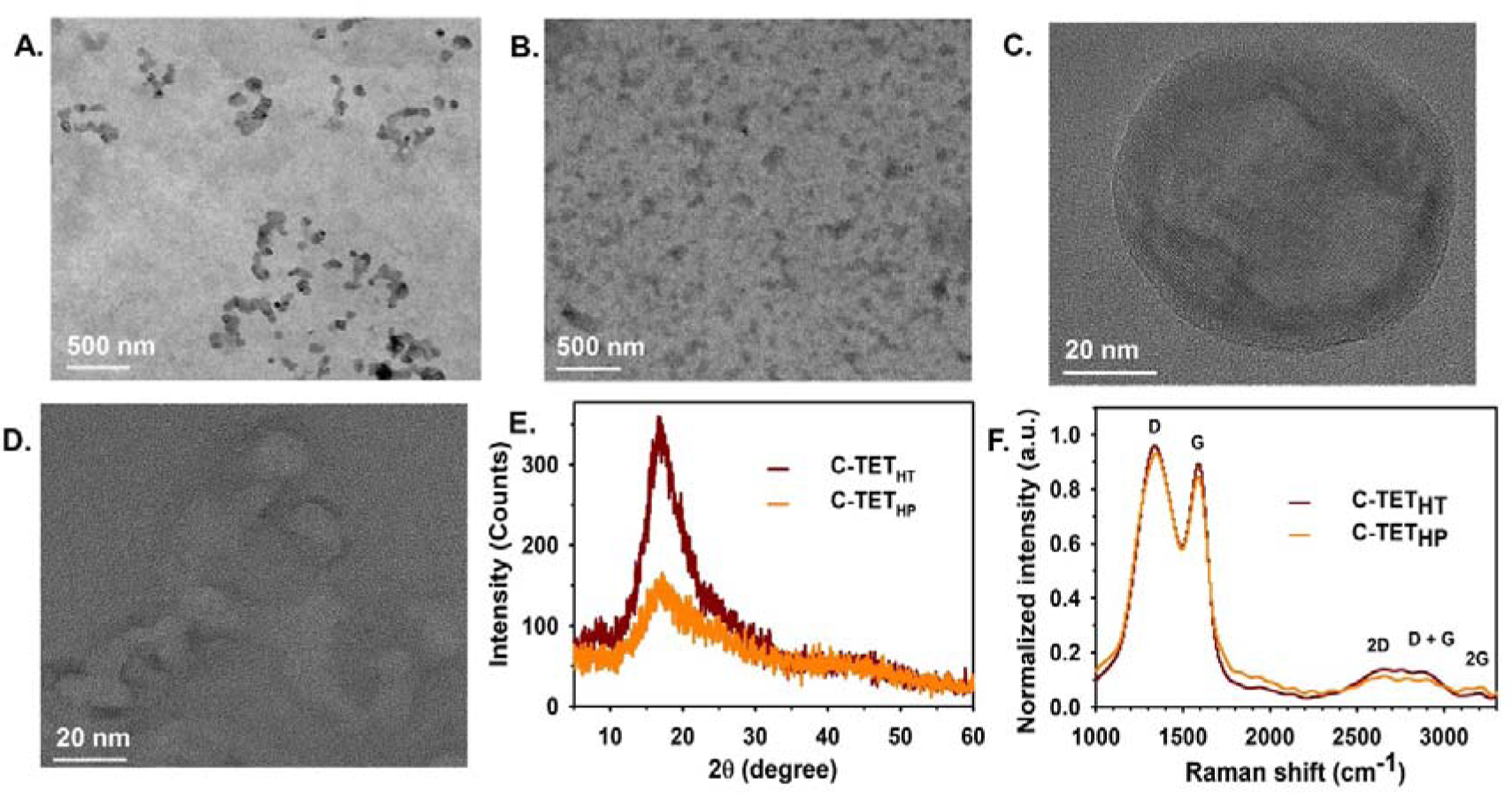
TEM image of **A.** C-TET_HT_, and **B.** C-TET_HP_. HR-TEM image of **C.** C-TET_HT_, and **D.** C-TET_HP_. The TEM imaging revealed a quasi-spherical morphology of C-TET_HT_ with a size of 70±26 nm, whereas C-TET_HP_ showed no definable structure. **E.** pXRD, and **F.** Raman spectra of C-TET_HT_ and C-TET_HP_ showed partial graphitization.

The EDS analysis showed the atomic composition (%) of the C-TETs as carbon 85.33±2.45, nitrogen 0.40±0.24, and oxygen 14.27±2.45 for C-TET_HT_ (**Figure S3**) and carbon 95.87±0.81, nitrogen 0.00±0.11, and oxygen 4.13±0.80 for C-TET_HP_ (**Figure S4**). As TET has an atomic composition (%) of carbon 68.75, nitrogen 6.25, and oxygen 25, it could be said that a significant amount of nitrogen and oxygen was lost during pyrolysis and the loss is less for the closely pyrolyzed C-TET_HT_ than the openly pyrolyzed C-TET_HP_. Sodium and chlorine which may have arisen in the sample either from the starting material or during the pH adjustment, were omitted from this calculation of atomic composition for ease of understanding and comparison with other analytical studies. The DLS analysis disclosed a much higher polydispersity index (PDI) for C-TET_HP_ (0.76±0.06) as compared to the PDI of C-TET_HT_ (0.33±0.00) indicating the superiority of hydrothermal pyrolysis over hot plate pyrolysis in getting particles with lower degree of non-uniformity. The zeta potential of the C-TET_HT_ and C-TET_HP_ was determined as −14.84±2.57 mV and −13.42±1.66 mV, respectively, indicating no major differences in the overall surface charge of both C-TETs (**Figure S5**). The surface chemistry of the C-TETs was investigated through XPS study.^33^ The atomic composition (%) of the C-TET_HT_ was found as carbon 71.95±1.33, nitrogen 4.09±0.79, and oxygen 23.97±0.54 and for the C-TET_HP_ the atomic composition (%) was carbon 77.72±1.97, nitrogen 2.66±0.81, and oxygen 19.62±2.26. Noticeably, the nitrogen and oxygen are more at the surface of the C-TETs as compared to the whole C-TETs structure for both the C-TET_HT_ and C-TET_HP_ (compare the atomic composition % from the EDS). To explore the nature of the atomic bonding on the surface of C-TETs (up to 10 nm depth), high-resolution XPS of carbon, nitrogen, and oxygen was performed (**Figure 3, Table S2**). The C1s high-resolution XPS spectra of C-TET_HT_ and C-TET_HP_ were deconvoluted to C=C/C-C, C-N/C-O, and C=O. Notably, a higher C-O/C-N content was detected for C-TET_HT_ (29.51 vs 11.87 for C-TET_HP_). The deconvoluted N1s high-resolution spectra unveiled the presence of a higher percentage of graphitic N in C-TET_HT_ (29.60) as compared to C-TET_HP_ (3.09). The O1s high-resolution spectra disclosed higher C-O content for C-TET_HT_ (60.88 vs 7.69 for C-TET_HP_). All these high-resolution XPS revelations indicate that C-TET_HT_ has a bigger interconnected structure, which is very substantial when we re-consider the visualized graphitic structures of the C-TETs in HR-TEM. Expectedly 1H NMR of the C-TETs revealed broad peaks in the aromatic, amide, alcohol, ketone, and alkyl regions indicating the presence of diverse types of chemical functionality in the C-TETs (**Figure S6**).^36^ Noticeably the carbonization process induced a decent amount of aromaticity in both the C-TETs.

**Figure 3.**
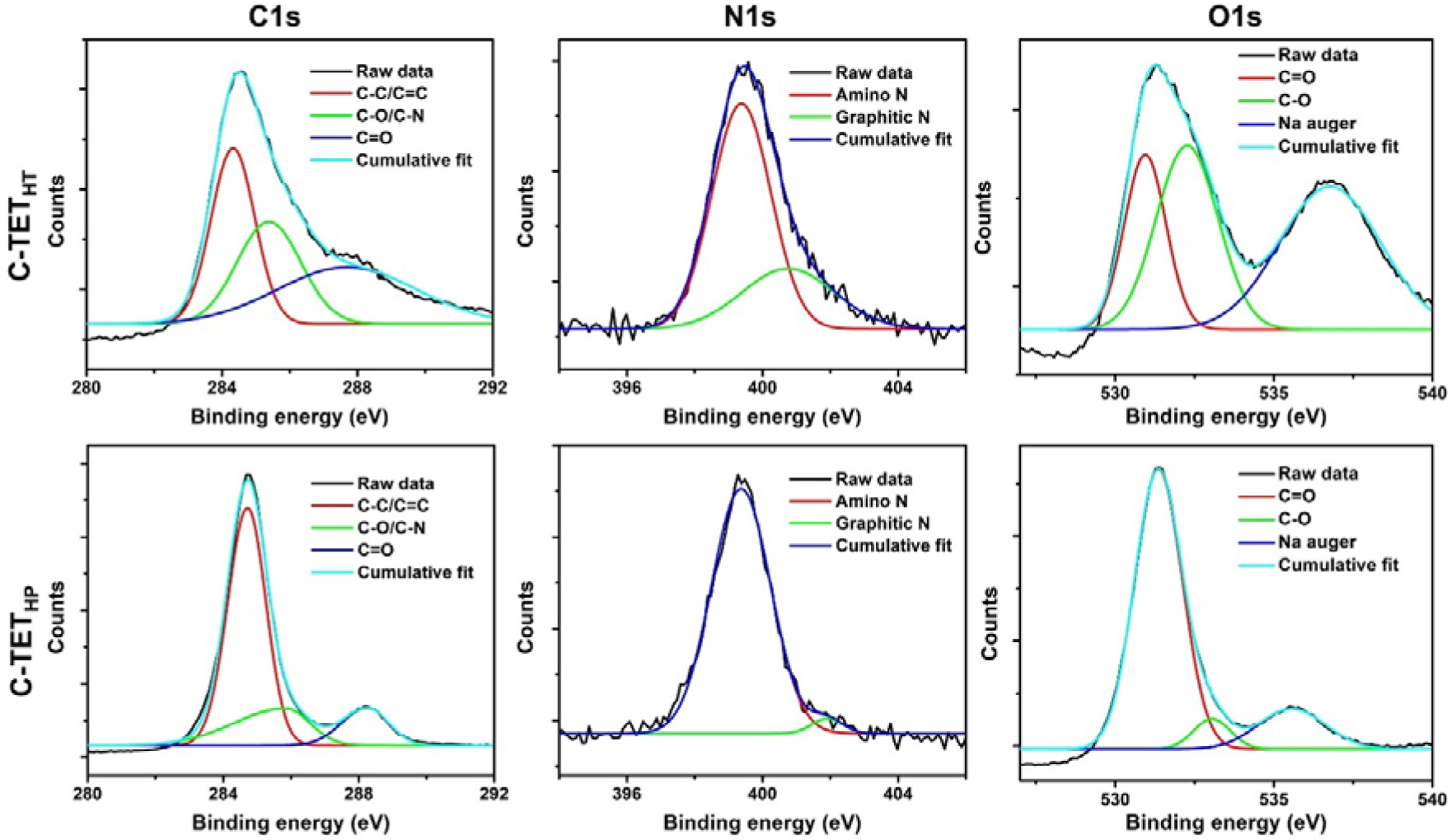
High-resolution XPS spectra of C-TET_HT_ and C-TET_HP_ unveiled the nature of the atomic bonding on the surface.

The photophysical behavior of the C-TETs was investigated through UV-Vis absorption and steady-state fluorescence spectroscopy. UV-Vis absorption spectra of the C-TETs showed no distinct peak, rather the absorption is spread and covered the whole visible region indicating the presence of several low-energy electronic transition states (**Figure S7A**).^33^ The steady-state fluorescence spectra of the C-TETs showed that the excitation-independent fluorescence emission is centered at ∼415 nm with excitation maxima at 325 nm (**Figure S7B-C**). Notably, this suggests fluorescence emission from uniform surface states,^3^ i.e., there is no significant difference in the surface state of the C-TET_HT_ and C-TET_HP_.

### 3.4. Photoinduced ROS generation and C-TETs

Photo-induced ROS production by photosensitizing drugs is the main reason for phototoxicity. There are reports that the tetracyclines class of drugs could induce phototoxicity on the skin.^37^ To examine whether the C-TETs could produce ROS in the presence of light, we performed the DHR 123 assay as per standard protocol.^30^ Neither the C-TETHT nor the C-TETHP have shown photoinduced ROS generation capacity (**Figure S8**). Instead, they protected the photo-oxidation of DHR 123 to rhodamine 123 another testimony of their antioxidant potential.

### 3.5. Cytotoxicity evaluation of C-TETs

The cytotoxicity of the C-TET_HT_ and C-TET_HP_ were evaluated on the NHLF cell line. The metabolic activity of the NHLF cells in the presence and absence of different concentrations of C-TETs and TET was measured with PrestoBlue assay (**Figure 4A, Table S3**). As compared to the no treatment, both the C-TET_HT_ and C-TET_HP_ showed no significant suppression in metabolic activity up to day 3 except the C-TET_HP_ at 100 µg/mL (p-value 0.005). However, as compared to the TET at 100 µg/mL on day 3 which showed substantial suppression in metabolic activity as compared to no treatment (p-value 0.001), this suppression in metabolic activity has no significance (p-value 0.863). Moreover, C-TET_HT_ at 100 µg/mL on day 3 showed better metabolic activity as compared to the TET at 100 µg/mL (p-value 0.048). The cell viability assay showed quite similar trends (**Figure 4B**). Excitingly, both the PrestoBlue assay and cell viability assay confirmed better biocompatibility of the C-TET_HT_ and C-TET_HP_ as compared to the TET. Additionally, the better biocompatibility of the C-TET_HT_ than the C-TET_HP_ specifies the importance of the carbonized parameters such as morphology, size, shape, and chemical functionality on cytotoxicity. Overall, these revelations encouraged us to study the non-antibiotic therapeutic potential of the C-TET_HT_ and C-TET_HP_ on other cell lines.

**Figure 4.**
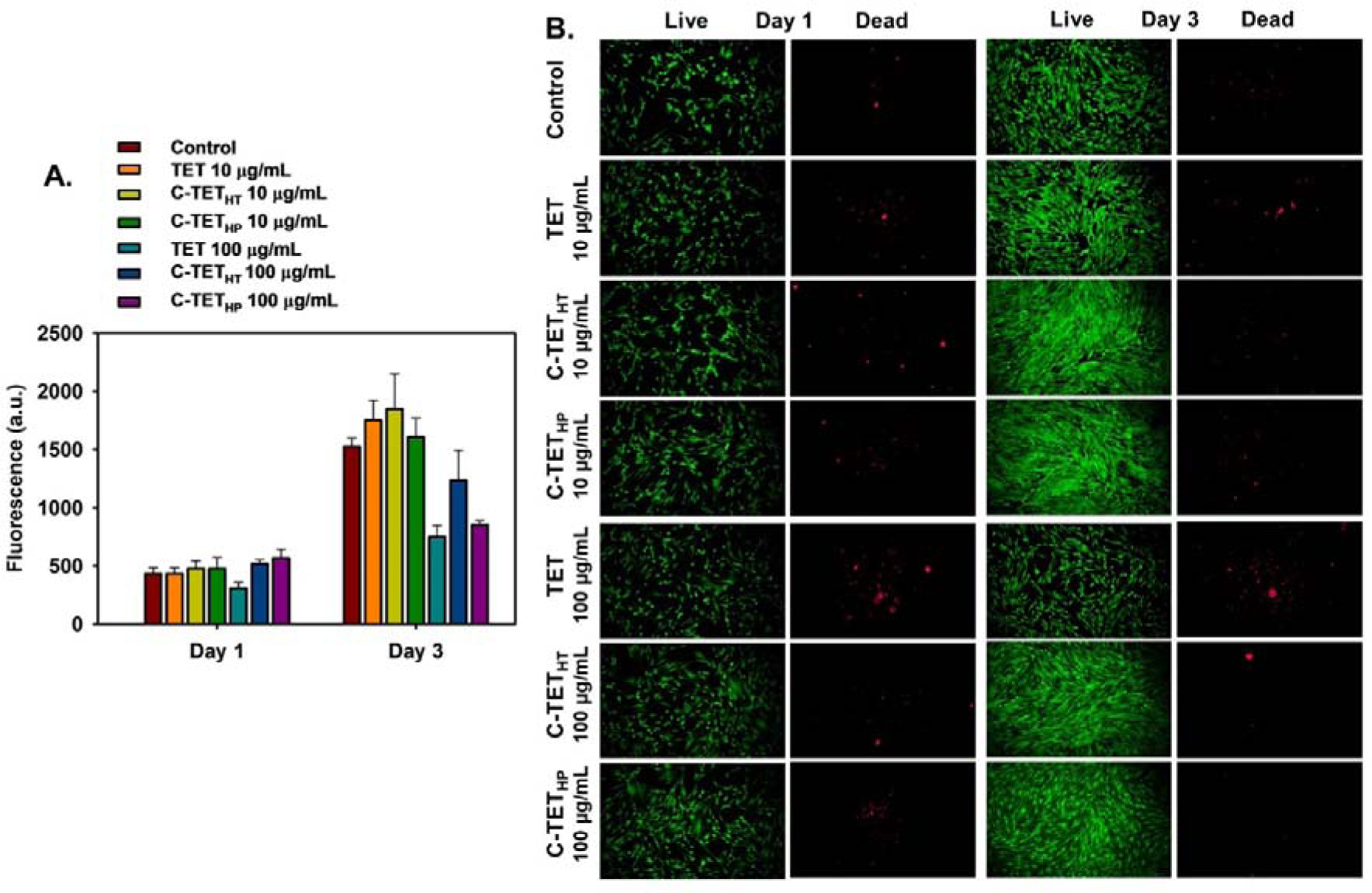
**A.** PrestoBlue assay and **B.** Cell viability assay of TET, C-TET_HT,_ and C-TET_HP_ showed lower cytotoxicity of the C-TETs as compared to TET.

### 3.6. Macrophage polarization by the C-TETs

The macrophage polarization potential of the C-TET_HT_ and C-TET_HP_ was evaluated through quantitative gene expression analysis. Both the C-TET_HT_ and C-TET_HP_ effectively modulated the phenotype of pro-inflammatory M1 macrophage to a pro-regenerative M2 phenotype by significantly downregulating the expression of *IL6* and *IL1b* in a dose-dependent manner (**Figure 5**). Whereas no significant phenotype changes were observed for M0 macrophages after the treatment with C-TET_HT_ and C-TET_HP_ (**Figure S9, Table S4**). Nevertheless, we have not observed notable correlations between the macrophage polarization and the carbonized parameters. Herein, we believe a study with a longer time course is needed to unveil some interesting correlations between the macrophage polarization and the carbonized parameters.

**Figure 5.**
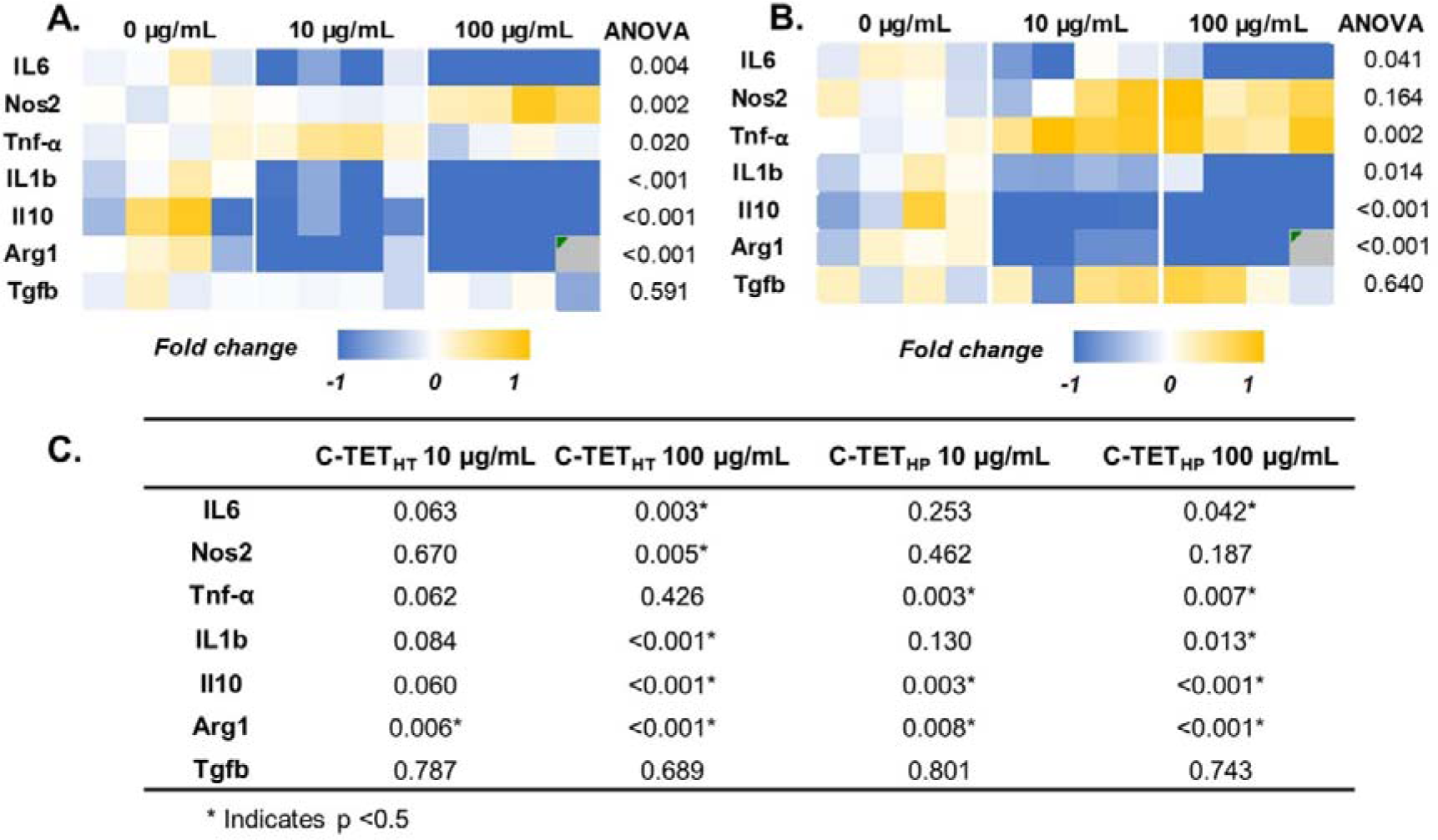
Heatmaps of the M1 macrophage gene expression after treatment with 10 µg/mL and 100 µg/mL of **A.** C-TET_HT_ and **B.** C-TET_HP_. Heatmaps show fold change with respect to the housekeeping gene (GAPDH) and untreated control. **C.** Corresponding Holm-Sidak pairwise comparison (p-value) with respect to untreated control.

### 3.7. Impact of the C-TETs on chondrogenesis

A drug or a material that can protect chondrocytes and promote chondrogenesis has great importance in osteoarthritis.^27^ Observing the ability of the C-TET_HT_ and C-TET_HP_ to modulate the macrophage phenotype of pro-inflammatory M1 macrophage to a pro-regenerative M2 phenotype, we become interested in exploring their potential to promote chondrogenesis under an inflammatory condition. Quantitative gene expression analysis after 1 day of treatment showed that C-TET_HT_ and C-TET_HP_ can attenuate the *IL1b*-induced pro-inflammatory responses by significantly downregulating the gene expression for COX2, ADAMTS 4 (aggrecanase 1), and MMP 9 (gelatinase) in a dose-dependent manner (**Figure 6**). C-TET_HP_ also downregulated the gene expression for MMP 3 (proteoglycanase) and MMP 13 (collagenase 3) in a dose-dependent manner. However, at a longer time course of treatment (Day 7) the ability of the C-TET_HT_ and C-TET_HP_ to attenuate the *IL1b*-induced pro-inflammatory responses mostly went away (**Figure S10, Table S5**). Even the viability of the ATDC5 cells was poor at higher concentrations (100 µg/mL) of the C-TET_HT_ and C-TET_HP_. The only positive outcome of the C-TET_HT_ and C-TET_HP_ 7-day treatment was the upregulation of COL2A1 and ACAN (aggrecan) gene expression at low (10 µg/mL) concentrations, which suggests criticality in selecting the dosage amount and time course. On the other hand, in terms of carbonized parameters, C-TET_HP_ appeared superior at low dosages.

**Figure 6.**
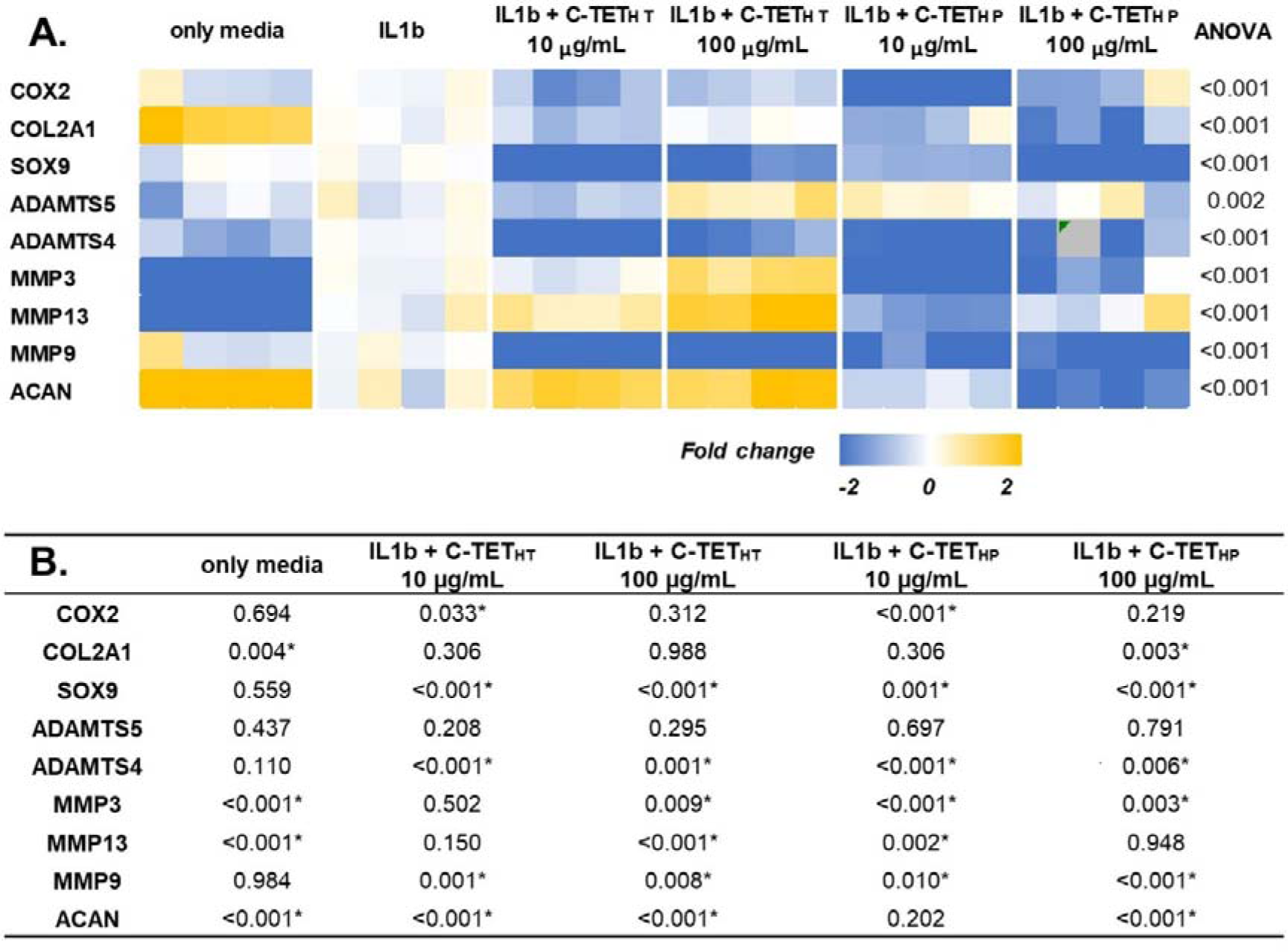
**A.** Heatmap of ATDC5 cells gene expression after 24 hr treatment with 10 µg/mL and 100 µg/mL of C-TET_HT_ and C-TET_HP_ in the presence of IL-1b. The heatmap shows fold change with respect to the housekeeping gene (GAPDH) and IL-1b treatment. B. Corresponding Holm-Sidak pairwise comparison (p-value) with respect to IL-1b treatment.

### 3.8. Osteogenic potential of the C-TETs

The osteogenic potential of the C-TET_HT_ and C-TET_HP_ was investigated by culturing the MC3T3-E1 cell line in normal growth media and osteogenic media for 3 weeks in the presence and absence of the C-TETs. Based on the previous observations of the low cell viability at a higher concentration of 100 µg/mL, here we studied 5 µg/mL as the low and 25 µg/mL as the high concentration of the C-TET_HT_ and C-TET_HP_. Quantitative gene analysis showed that osteogenic genes^32^ such as runt-related transcription factor 2 (Runx2), collagen type I alpha 1 (Col1a1), alkaline phosphatase 1 (Alp 1), integrin binding sialoprotein (Ibsp), and osterix (Sp7) were upregulated significantly by the C-TET_HT_ and C-TET_HP_ treatment in the normal media in a dosage-dependent manner (**Figure 7A, Table S6**). However, the ALP activity showed a substantial increase only for the C-TET_HP_ treatment at low concentrations (**Figure 7B, Figure S11, Table S7**). Whereas quantification of the mineralization (alizarin red staining) showed no significant differences among the groups (**Figure 7C, Figure S12, Table S8**). In the osteogenic media, gene expressions for Alp1, bone gamma-carboxyglutamate protein (Bglap, osteocalcin), osteopontin (Spp1), Ibsp, and Sp7 were upregulated significantly in a dosage-dependent manner by the C-TET_HT_ and C-TET_HP_ (**Figure 8A, Table S9**). The ALP activity assay and alizarin red staining did not show a significant increase in ALP activity (**Figure 8B, Table S10**) and mineralization (**Figure 8C, Figure S13, Table S11**) as compared to the untreated control in the osteogenic media. These contrasting results among the gene expression and ALP activity assay and mineralization are due to the poor cell viability at higher concentrations of the C-TETs. However, the possibility of hindered ALP activity and mineralization in the presence of the C-TETs cannot be excluded.^38^ Nevertheless, through this study C-TET_HP_ has emerged as a superior osteogenic nanomaterial at low concentration that again highlights the importance of carbonized parameters on cellular fates.

**Figure 7.**
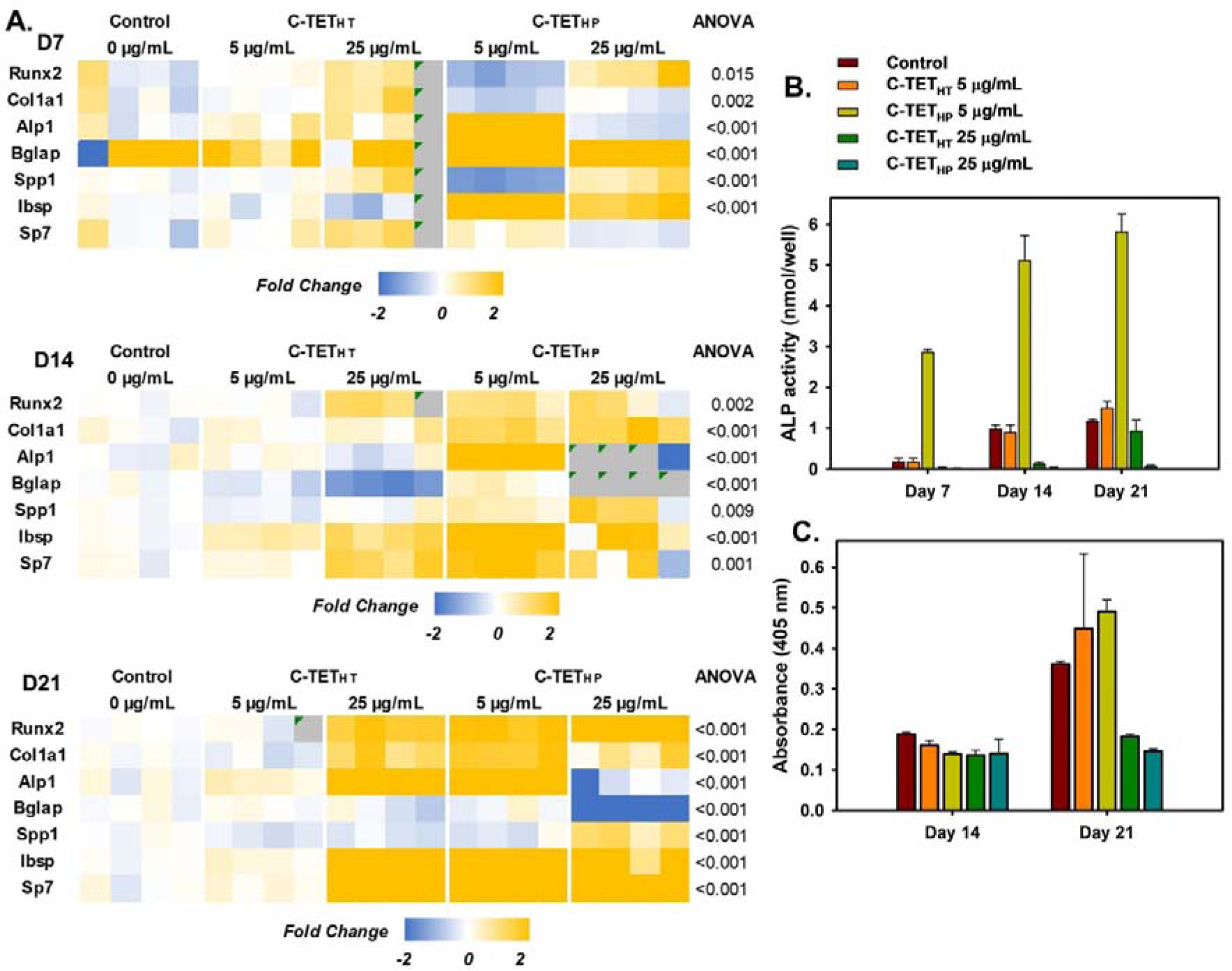
**A.** Heatmaps of MC3T3-E1 gene expression on different time courses treated with C-TET_HT_ and C-TET_HP_ in normal growth media. Heatmaps show fold change with respect to the housekeeping gene (GAPDH) and untreated control. **B.** ALP activity, and **C.** quantification of mineralization (alizarin red staining) of MC3T3-E1 cells on different time courses.

**Figure 8.**
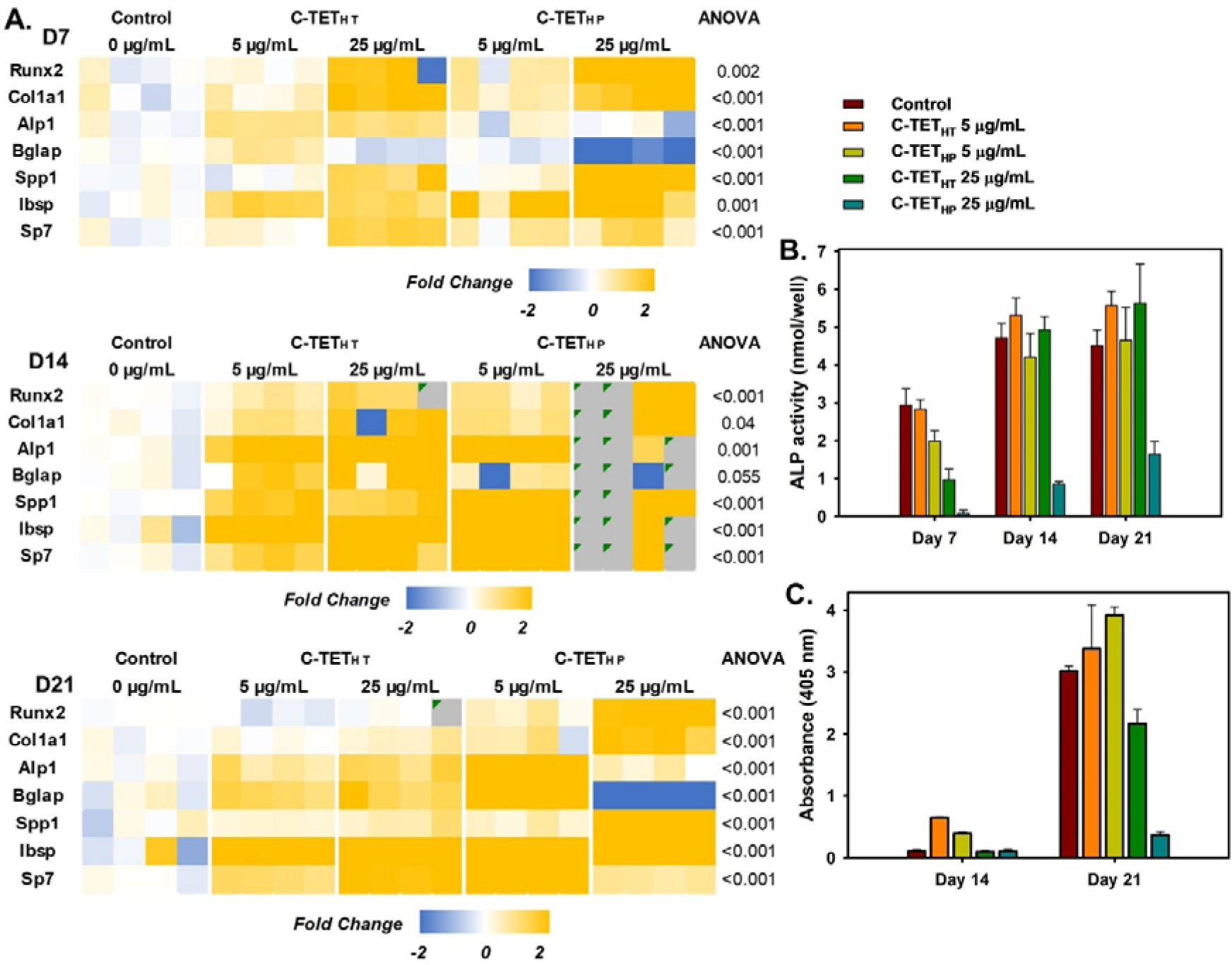
**A.** Heatmaps of MC3T3-E1 gene expression on different time courses treated with C-TET_HT_ and C-TET_HP_ in osteogenic growth media. Heatmaps show fold change with respect to the housekeeping gene (GAPDH) and untreated control. **B.** ALP activity, and **C.** quantification of alizarin red staining of MC3T3-E1 cells on different time course.

## 4. Conclusion

Enhancing the overall efficacy of the therapeutic potential of a drug is constructed with a series of rigorous activities that start with computational studies, organic synthesis, derivatization, etc. Herein, in an exclusive approach, we have synthesized a series of carbonized-Tetracyclines (C-TETs) to reduce the toxicity or adverse effects of TET while retaining or improving a few of its non-antibiotic properties. After varying a series of carbonization conditions in terms of solvent pH, method of pyrolysis, temperature, and time, we got a library of C-TETs with good water dispersibility, i.e., partial carbonization can improve the water solubility of a poorly water-soluble drug. Strikingly, the antioxidant capacity of those C-TETs was found 10-20 better than that of the TET. For detailed characterization and evaluation of a few non-antibiotic properties, we have shorted out two C-TETs, C-TET_HT_ and C-TET_HP_, where the sole difference in their synthesis parameters is in the method of pyrolysis. C-TET_HT_ was synthesized in a closed hydrothermal apparatus, while the C-TET_HP_ was synthesized by heating openly on a hot plate. The analytical characterization of the C-TET_HT_ and C-TET_HP_ have shown distinct differences in morphology, size, shape, and chemical functionality. Where the C-TET_HT_ appeared as a comparatively bulk (70±26, round-shaped) graphitic structure, the C-TET_HP_ is made of tiny (<10 nm) carbon fragments with no definable structure. Additionally, nitrogen and oxygen content were found more in C-TET_HT_ than the C-TET_HP_. Both the C-TETs have no ROS generation capacity in the presence of light nullifying the chance of phototoxicity associated with the tetracyclines class of drugs. Remarkably, both the C-TET_HT_ and C-TET_HP_ appeared less cytotoxic than the TET, where the C-TET_HT_ showed the least cytotoxicity. Both the C-TETs modulate the pro-inflammatory M1 macrophage to pro-regenerative M2 macrophage in a dose-dependent manner. Although treatment with the C-TETs initially mitigated the pro-inflammatory effect of *IL1b*, their long-time use has been questioned. Through osteogenicity studies, the C-TET_HP_ has emerged as a decent osteogenic nanomaterial in a dose-dependent manner. Thus, we have found good correlations between different non-antibiotic properties of the C-TETs and their carbonized forms in terms of morphology, size, shape, and chemical functionality. Furthermore, the dosage amount and exposure times appeared critical to the efficacy of both the C-TET_HT_ and C-TET_HP_. Although a lot of research needs to be performed before any concluding remark on the therapeutic potential of the C-TETs, overall, the C-TETs have shown substantial initial promises for further exploration in different musculoskeletal diseases. We believe our approach of nano-carbonization of small drug molecules to improve the overall therapeutic efficacy has a long way to go subject to rigorous research explorations.

## 5. Data availability statement

The original contributions presented in the study are included in the article/Supplementary Material. Further inquiries can be directed to the corresponding author.

## 6. Ethics statement

All studies involving animal or human cells are conducted according to approved protocols by the Institutional Biosafety Committee (IBC) protocol at the University of Kentucky.

## Acknowledgments

Research reported in this publication was supported in part by the National Institute of Arthritis and Musculoskeletal and Skin Diseases (NIAMS) Award Number R21AR078447, National Institute of General Medical Sciences (NIGMS) of the National Institutes of Health under Award Number P20GM130456, Orthopedic Trauma Association (OTA, Grant Number: 6889). The content is solely the responsibility of the authors and does not necessarily represent the official views of the National Institutes of Health or other grant funding agencies. SM thanks Matthew D. Patrick for helping with cell cultures and statistical analysis.

## 8. Supplementary Information

Additional data related to this work can be found on the supplementary information pages.

