## supplementary information for "Carbonized Tetracycline: a new class of nanomaterial with tuneable antioxidant, reduced cytotoxicity, immunomodulatory, and osteogenic properties"

University at Buffalo,

Buffalo, NY 14260

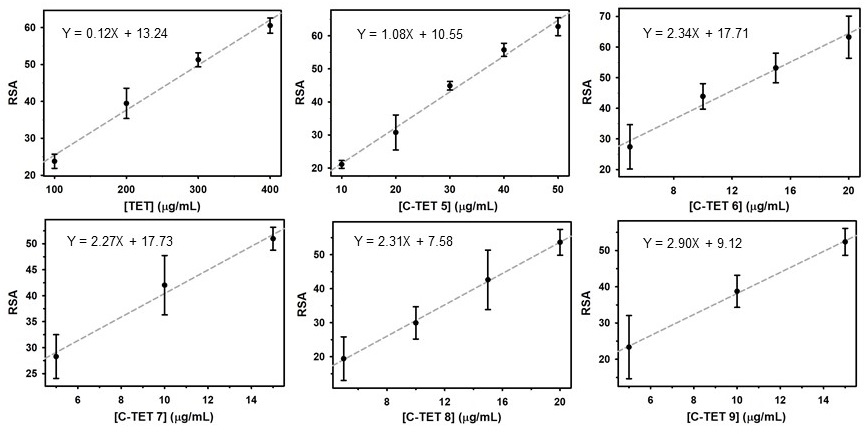

**Figure S1**. Standard curve for radical scavenging activity (RSA) vs concentration of TET and C-TETs.

**Table S1.** Calculation of the antioxidant value (IC_50_) of C-TETs from the standard curve.

| **C-TETs** | **Slope** | **Intercept** | **R^2^** | **IC_50_, µg/mL** |
| --- | --- | --- | --- | --- |
| TET | 0.12 | 13.24 | 0.99 | 301.08 |
| C-TET 5 | 1.08 | 10.55 | 0.99 | 36.42 |
| C-TET 6 | 2.34 | 17.71 | 0.98 | 13.81 |
| C-TET 7 | 2.27 | 17.73 | 0.99 | 14.22 |
| C-TET 8 | 2.31 | 7.58 | 1 | 18.40 |
| C-TET 9 | 2.90 | 9.12 | 1 | 14.08 |

R^2^ is the coefficient of determination.

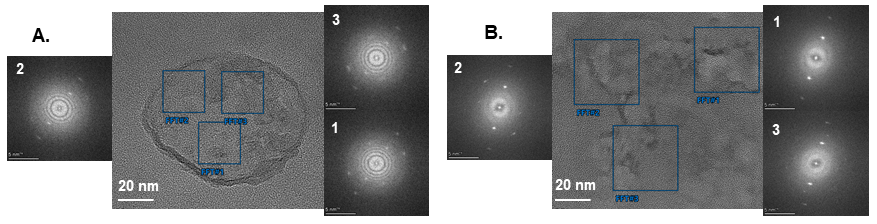

**Figure S2.** HR-TEM and the corresponding FFT of **A.** C-TET_HT_ and **B.** C-TET_HP_.

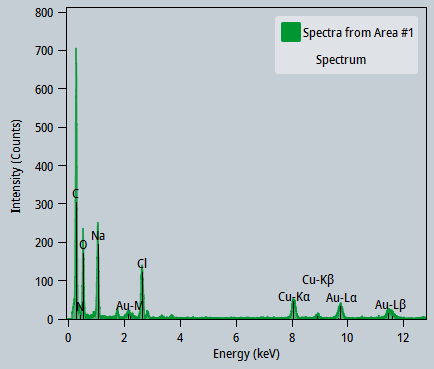

**Figure S3.** EDS spectra of C-TET_HT_.

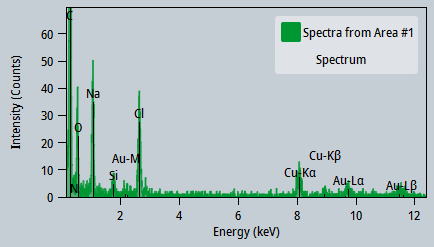

**Figure S4.** EDS spectra of C-TET_HP_.

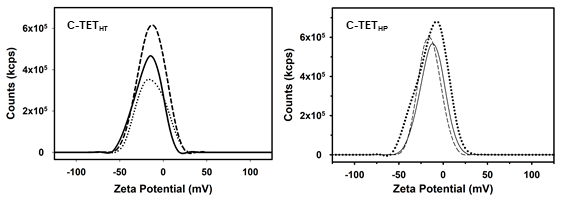

**Figure S5.** Zeta potential of C-TET_HT_ and C-TET_HP_.

**Table S2.** Analysis of high-resolution XPS spectra (C, N, and O) of the C-TET_HT_ and C-TET_HP_ through deconvolution.

| C-TET_HT_ | C1s | C-C/C=C | 34.71 |
| --- | --- | --- | --- |
|  |  | C-O/C-N | 29.51 |
|  |  | C=O | 35.77 |
|  | N1s | Amino N | 70.40 |
|  |  | Graphitic N | 29.60 |
|  | O1s | C=O | 39.12 |
|  |  | C-O | 60.88 |
| C-TET_HP_ | C1s | C-C/C=C | 76.06 |
|  |  | C-O/C-N | 11.87 |
|  |  | C=O | 12.08 |
|  | N1s | Amino N | 96.91 |
|  |  | Graphitic N | 3.09 |
|  | O1s | C=O | 92.31 |
|  |  | C-O | 7.69 |

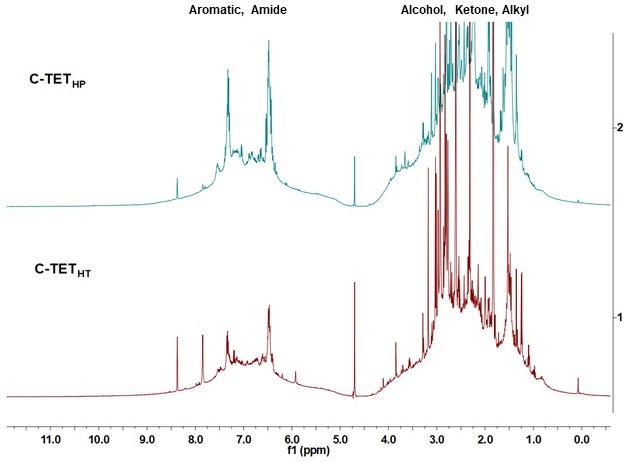

**Figure S6.** ^1^H NMR of C-TET_HT_ and C-TET_HP_.

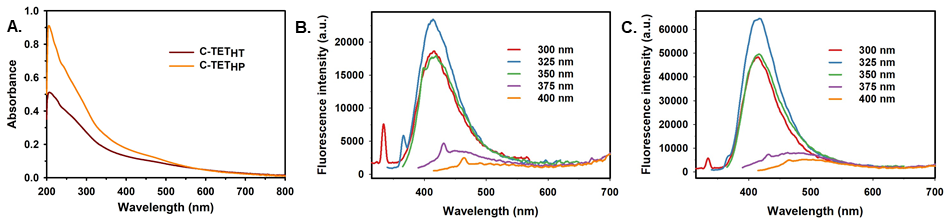

**Figure S7.** **A.** UV-Vis absorption spectra, and **B-C.** steady-state fluorescence spectra of C-TET_HT_ and C-TET_HP_.

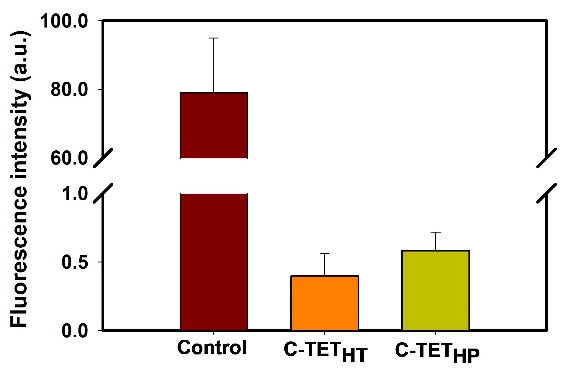

**Figure S8.** ROS generation study of the C-TET_HT_ and C-TET_HP_ using DHR 123 as a probe.

**Table S3.** Holm-Sidak comparisons (p-value) for PrestoBlue assay.

|  | **Day 1** | | |  | **Day 3** | |
| --- | --- | --- | --- | --- | --- | --- |
|  | **Control** | **TET, 10 µg/mL** | **TET, 100 µg/mL** | **Control** | **TET, 10 µg/mL** | **TET, 100 µg/mL** |
| **Control** |  | 1.000 | 0.229 |  | 0.522 | 0.001* |
| **C-TET_HT_ 10 µg/mL** | 0.961 | 0.961 |  | 0.284 | 0.863 |  |
| **C-TET_HT_ 100 µg/mL** | 0.593 |  | 0.001* | 0.362 |  | 0.048* |
| **C-TET_HP_ 10 µg/mL** | 0.961 | 0.961 |  | 0.863 | 0.800 |  |
| **C-TET_HP_ 100 µg/mL** | 0.156 |  | 0.008* | 0.005* |  | 0.863 |

**B.**

**A.**

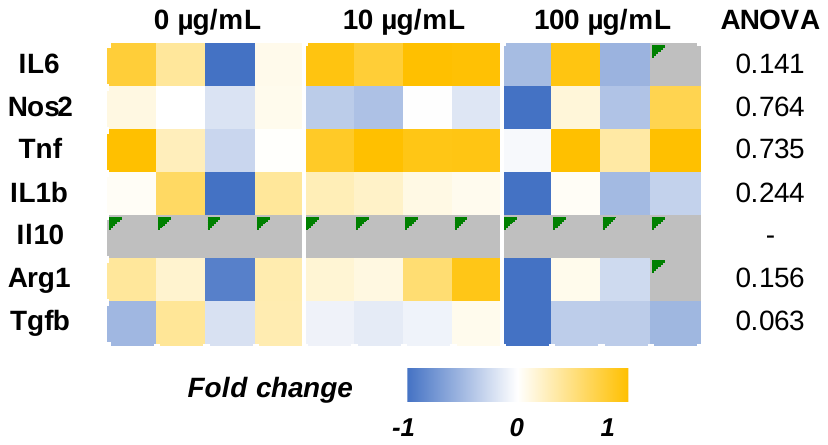

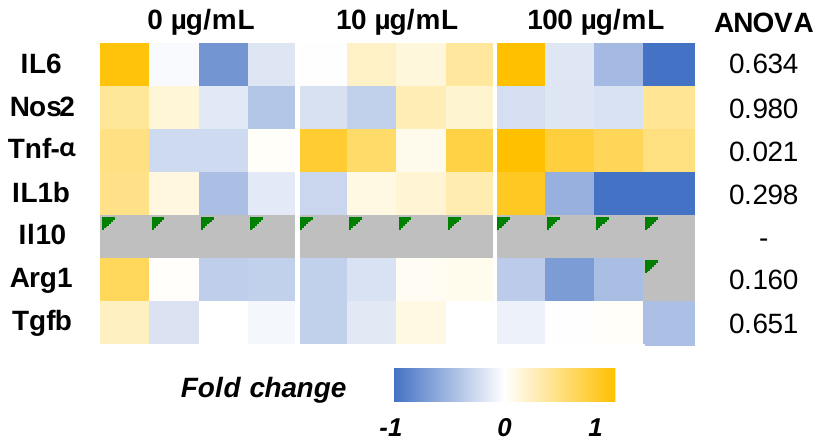

**Figure S9.** Heatmaps of the M0 macrophage gene expression after treatment with 10 µg/mL and 100 µg/mL of **A.** C-TET_HT_ and **B.** C-TET_HP_. Heatmaps show fold change with respect to the housekeeping gene (GAPDH) and untreated control.

**Table S4.** Holm-Sidak comparisons (p-value) with respect to untreated control for M0 macrophage.

| **Genes** | **C-TET_HT_ 10 µg/mL** | **C-TET_HT_ 100 µg/mL** | **C-TET_HP_ 10 µg/mL** | **C-TET_HP_ 100 µg/mL** |
| --- | --- | --- | --- | --- |
| **IL6** | 0.231 | 0.964 | 0.785 | 0.785 |
| **Nos2** | 0.876 | 0.876 | 0.996 | 0.996 |
| **Tnf-α** | 0.840 | 0.840 | 0.067 | 0.024* |
| **IL1b** | 0.644 | 0.410 | 0.883 | 0.417 |
| **Il10** | - | - | - | - |
| **Arg1** | 0.482 | 0.482 | 0.662 | 0.199 |
| **Tgfb** | 0.807 | 0.097 | 0.774 | 0.764 |

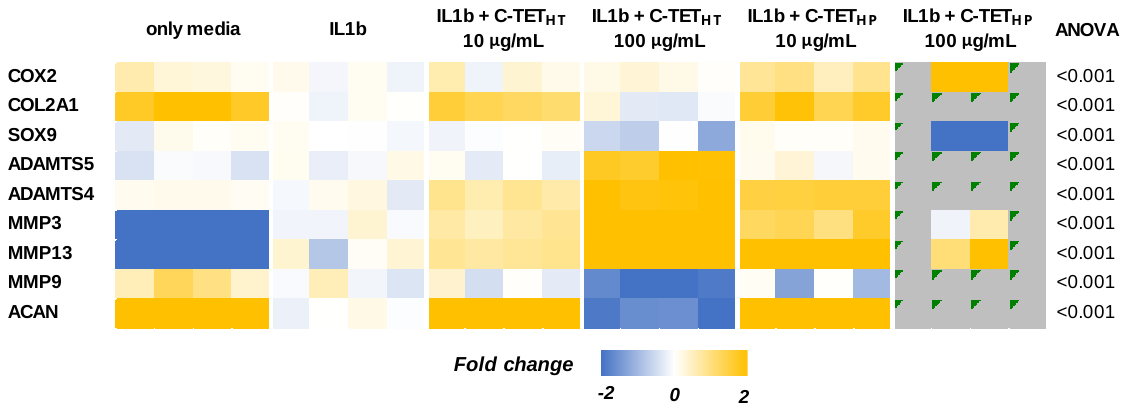

**Figure S10.** Heatmap of ATDC5 cells gene expression after 7 days of treatment with 10 µg/mL and 100 µg/mL of C-TET_HT_ and C-TET_HP_ in the presence of IL-1b. The heatmap shows fold change with respect to the housekeeping gene (GAPDH) and IL-1b treatment.

**Table S5.** Holm-Sidak pairwise comparison (p-value) with respect to IL-1b treatment for 7 days on ATDC5 cells (gene expression).

|  | **only media** | **IL1b + C-TET_HT_ 10 µg/mL** | **IL1b + C-TET_HT_ 100 µg/mL** | **IL1b + C-TET_HP_ 10 µg/mL** | **IL1b + C-TET_HP_ 100 µg/mL** |
| --- | --- | --- | --- | --- | --- |
| **COX2** | 0.225 | 0.460 | 0.660 | <0.001* | <0.001* |
| **COL2A1** | <0.001* | <0.001* | 0.606 | <0.001* | - |
| **SOX9** | 0.999 | 0.999 | 0.126 | 0.997 | <0.001* |
| **ADAMTS5** | 0.450 | 0.707 | <0.001* | 0.707 | - |
| **ADAMTS4** | 0.166 | <0.001* | <0.001* | <0.001* | - |
| **MMP3** | <0.001* | 0.040* | <0.001* | <0.001* | 0.419 |
| **MMP13** | <0.001* | 0.031* | <0.001* | <0.001* | <0.001* |
| **MMP9** | 0.172 | 0.828 | <0.001* | 0.419 | - |
| **ACAN** | <0.001* | <0.001* | <0.001* | <0.001* | - |

**Table S6.** Holm-Sidak pairwise comparison (p-value) of MC3T3-E1 gene expression on different time courses treated with C-TET_HT_ and C-TET_HP_ in normal growth media.

|  | **D7** | | | | **D14** | | | | **D21** | | | |
| --- | --- | --- | --- | --- | --- | --- | --- | --- | --- | --- | --- | --- |
|  | **C-TET_HT_** | | **C-TET_HP_** | | **C-TET_HT_** | | **C-TET_HP_** | | **C-TET_HT_** | | **C-TET_HP_** | |
|  | **5 µg/mL** | **25 µg/mL** | **5 µg/mL** | **25 µg/mL** | **5 µg/mL** | **25 µg/mL** | **5 µg/mL** | **25 µg/mL** | **5 µg/mL** | **25 µg/mL** | **5 µg/mL** | **25 µg/mL** |
| **Runx2** | 0.889 | 0.646 | 0.562 | 0.188 | 0.859 | 0.013* | 0.037* | 0.183 | 0.931 | <0.001* | <0.001* | <0.001* |
| **Col1a1** | 0.947 | 0.034* | 0.301 | 0.934 | 0.516 | 0.313 | <0.001* | <0.001* | 0.338 | 0.001* | <0.001* | 0.032* |
| **Alp1** | 0.303 | 0.263 | <0.001* | 0.303 | 0.721 | 0.721 | <0.001* | <0.001* | 0.341 | <0.001* | <0.001* | 0.102 |
| **Bglap** | 0.626 | 0.626 | <0.001* | 0.018* | 0.071 | <0.001* | 0.071 | - | 0.999 | 0.999 | 0.999 | <0.001* |
| **Spp1** | 0.991 | 0.012* | <0.001* | 0.011* | 0.737 | 0.737 | 0.519 | 0.041* | 0.345 | 0.345 | 0.463 | <0.001* |
| **Ibsp** | 0.988 | 0.062 | <0.001* | <0.001* | 0.478 | 0.066 | <0.001* | 0.066 | 0.547 | <0.001* | <0.001* | <0.001* |
| **Sp7** | 0.678 | 0.03* | 0.534 | 0.678 | 0.715 | 0.045* | 0.003* | 0.576 | 0.477 | <0.001* | <0.001* | <0.001* |

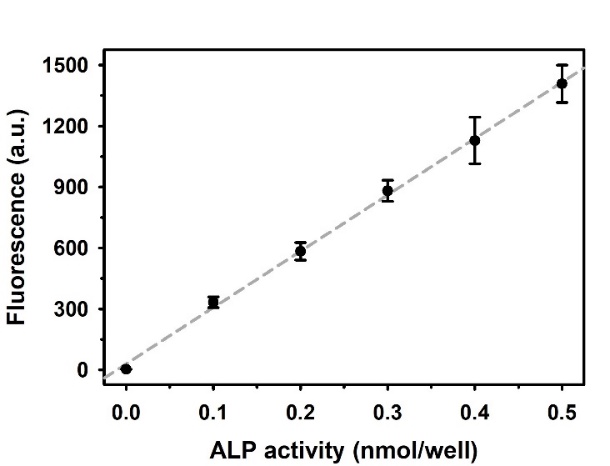

**Figure S11.** Standard curve of ALP activity.

**Table S7.** Holm-Sidak pairwise comparison (p-value) with respect to untreated control for ALP activity of the MC3T3-E1 cells grown in normal media.

|  | **Day 7 (P <0.001)** | **Day 14 (P <0.001)** | **Day 21 (P <0.001)** |
| --- | --- | --- | --- |
| **C-TET_HT_ 5 µg/mL** | 0.808 | 0.851 | 0.148 |
| **C-TET_HT_ 25 µg/mL** | 0.002* | 0.003* | 0.167 |
| **C-TET_HP_ 5 µg/mL** | <0.001* | <0.001* | <0.001* |
| **C-TET_HP_ 25 µg/mL** | <0.001* | 0.002* | <0.001* |

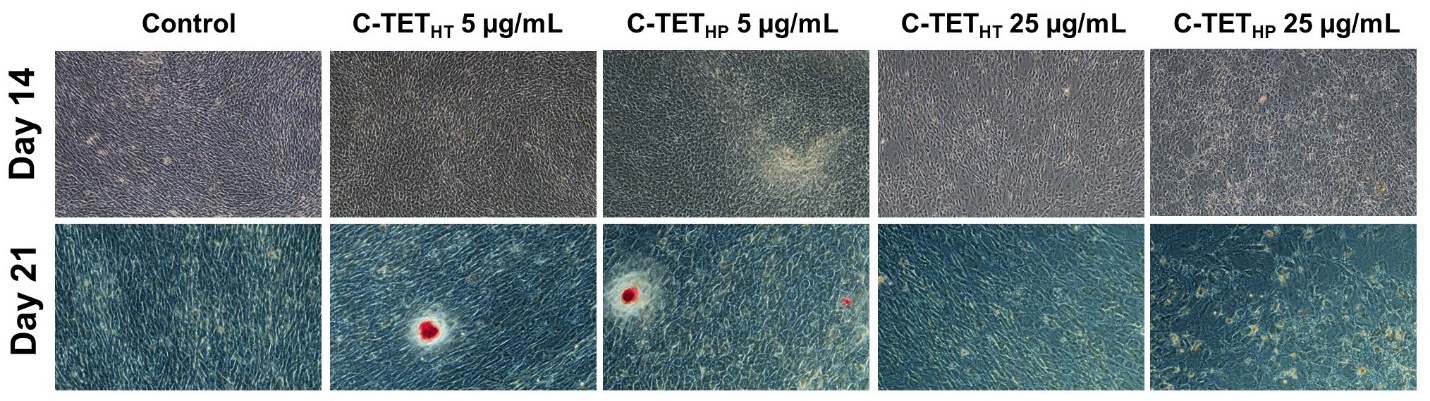
**Figure S12.** Bright-field image of alizarin red staining of MC3T3-E1 cells grown in normal media.

**Table S8.** Holm-Sidak pairwise comparison (p-value) with respect to untreated control for alizarin red staining of the MC3T3-E1 cells grown in normal media.

|  | **Day 14 (P <0.091)** | **Day 21 (P <0.028)** |
| --- | --- | --- |
| **C-TET_HT_ 5 µg/mL** | 0.674 | 0.720 |
| **C-TET_HT_ 25 µg/mL** | 0.209 | 0.353 |
| **C-TET_HP_ 5 µg/mL** | 0.229 | 0.552 |
| **C-TET_HP_ 25 µg/mL** | 0.229 | 0.261 |

**Table S9.** Holm-Sidak pairwise comparison (p-value) of MC3T3-E1 gene expression on different time courses treated with C-TET_HT_ and C-TET_HP_ in osteogenic growth media.

|  | **D7** | | | | **D14** | | | | **D21** | | | |
| --- | --- | --- | --- | --- | --- | --- | --- | --- | --- | --- | --- | --- |
|  | **C-TET_HT_** | | **C-TET_HP_** | | **C-TET_HT_** | | **C-TET_HP_** | | **C-TET_HT_** | | **C-TET_HP_** | |
|  | **5 µg/mL** | **25 µg/mL** | **5 µg/mL** | **25 µg/mL** | **5 µg/mL** | **25 µg/mL** | **5 µg/mL** | **25 µg/mL** | **5 µg/mL** | **25 µg/mL** | **5 µg/mL** | **25 µg/mL** |
| **Runx2** | 0.912 | 0.685 | 0.912 | 0.003 | 0.181 | 0.007 | 0.067 | <0.001 | 0.453 | 0.933 | 0.077 | <0.001 |
| **Col1a1** | 0.318 | <0.001 | 0.086 | <0.001 | 0.878 | 0.878 | 0.835 | 0.029 | 0.844 | 0.339 | 0.36 | <0.001 |
| **Alp1** | 0.023 | 0.015 | 0.922 | 0.749 | <0.001 | <0.001 | <0.001 | 0.001 | 0.007 | <0.001 | <0.001 | 0.189 |
| **Bglap** | 0.345 | 0.726 | 0.843 | <0.001 | 0.768 | 0.671 | 0.768 | 0.542 | 0.001 | <0.001 | <0.001 | <0.001 |
| **Spp1** | 0.945 | 0.006 | 0.885 | <0.001 | <0.001 | <0.001 | <0.001 | <0.001 | 0.442 | 0.051 | 0.279 | <0.001 |
| **Ibsp** | 0.087 | 0.114 | 0.021 | <0.001 | <0.001 | <0.001 | <0.001 | 0.032 | <0.001 | <0.001 | <0.001 | <0.001 |
| **Sp7** | 0.374 | <0.001 | 0.117 | 0.012 | 0.002 | <0.001 | <0.001 | <0.001 | <0.001 | <0.001 | <0.001 | <0.001 |

**Table S10.** Holm-Sidak pairwise comparison (p-value) with respect to untreated control for ALP activity of the MC3T3-E1 cells grown in osteogenic media.

|  | **Day 7 (P <0.001)** | **Day 14 (P <0.001)** | **Day 21 (P <0.001)** |
| --- | --- | --- | --- |
| **C-TET_HT_ 5 µg/mL** | 0.551 | 0.206 | 0.176 |
| **C-TET_HT_ 25 µg/mL** | 0.002 | 0.460 | 0.163 |
| **C-TET_HP_ 5 µg/mL** | <0.001 | 0.266 | 0.941 |
| **C-TET_HP_ 25 µg/mL** | <0.001 | <0.001 | <0.001 |

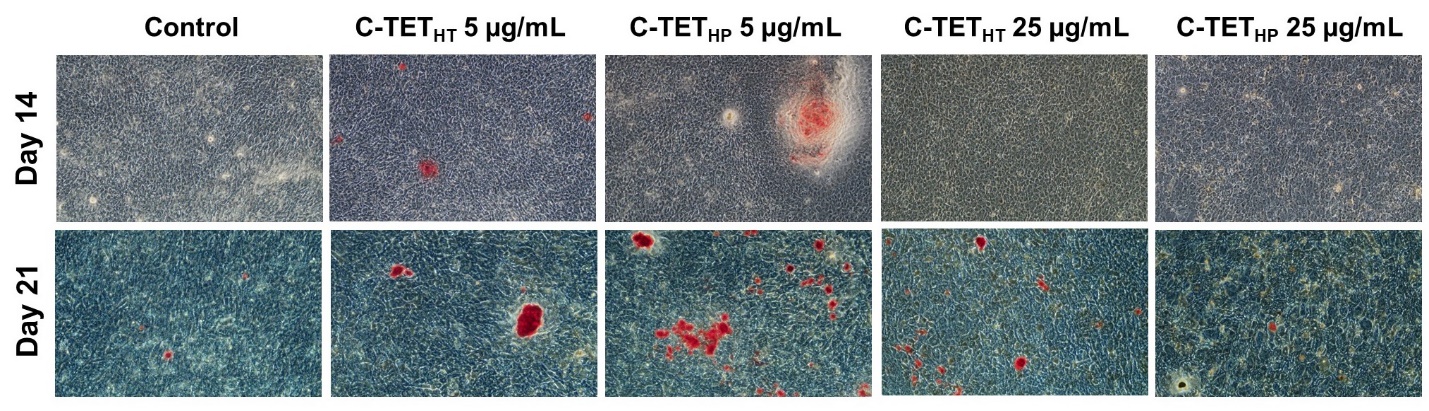
**Figure S13.** Bright-field image of alizarin red staining of MC3T3-E1 cells grown in osteogenic media.

**Table S11.** Holm-Sidak pairwise comparison (p-value) with respect to untreated control for alizarin red staining of the MC3T3-E1 cells grown in osteogenic media.

|  | **Day 14 (P <0.001)** | **Day 21 (P <0.001)** |
| --- | --- | --- |
| **C-TET_HT_ 5 µg/mL** | <0.001 | 0.337 |
| **C-TET_HT_ 25 µg/mL** | 0.863 | 0.163 |
| **C-TET_HP_ 5 µg/mL** | <0.001 | 0.163 |
| **C-TET_HP_ 25 µg/mL** | 0.863 | 0.004 |
